# Proximity Labeling-assisted Identification of Endogenous Kinase Substrates

**DOI:** 10.1101/2020.06.09.143370

**Authors:** Tomoya Niinae, Koshi Imami, Naoyuki Sugiyama, Yasushi Ishihama

**Affiliations:** Department of Molecular & Cellular BioAnalysis, Graduate School of Pharmaceutical Sciences, Kyoto University, 46–29 Yoshidashimoadachi-cho, Sakyo-ku, Kyoto 606–8501, Japan; PRESTO, Japan Science and Technology Agency (JST), 5-3 Yonban-cho, Chiyoda-ku, Tokyo, 102-0075, Japan

**Keywords:** Kinase substrate, BioID (proximity-dependent biotin identification), phosphoproteomics

## Abstract

Mass spectrometry-based phosphoproteomics can identify more than 10,000 phosphorylated sites in a single experiment. But, despite the fact that enormous phosphosite information has been accumulated in public repositories, protein kinase-substrate relationships remain largely unknown. Here, we describe a method to identify endogenous substrates of kinases by means of proximity labeling. We used a proximity-dependent biotin identification approach, called BioID, in combination with kinase-perturbed phosphoproteomics profiling and phosphorylation sequence motifs derived from *in vitro* kinase assay to find molecules that interact with a target kinase, that show altered phosphorylation in response to kinase perturbation, and that are directly phosphorylated by the kinase *in vitro*; i.e., endogenous kinase substrates. Application of this methodology to casein kinase 2 (CK2) and protein kinase A (PKA) identified 33 and 52 putative substrates, respectively. We also show that known cancer-associated missense mutations near phosphosites of substrates affect phosphorylation by CK2 or PKA, and thus might alter downstream signaling in cancer cells bearing these mutations.

This study extends our knowledge of kinase-substrate networks by proposing a new large-scale approach to identify endogenous substrates of kinases.

## Introduction

Protein phosphorylation plays a key role in intracellular signal transduction and regulates various biological processes, including cell proliferation and differentiation. Mass spectrometry (MS)-based phosphoproteomics has allowed us to identify thousands of phosphorylated sites in single experiments (1–3). However, despite the fact that enormous phosphosite information has been accumulated in public repositories (4–7), protein kinase-substrate relationships remain largely unknown both *in vitro* and *in vivo* (8).

*In vitro* kinase assay is one of the most widely used approaches to identify kinase substrates (9–13). We recently reported a total of 175,574 substrates for 385 kinases using *in vitro* kinase reaction with protein extracted from human cells, followed by phosphopeptide enrichment and liquid chromatography-tandem mass spectrometry (LC/MS/MS) analyses (14, 15). While this method successfully identified *in vitro* substrates and uncovered a variety of consensus motifs for kinase substrates, we still lack information on endogenous kinase substrates, as the physiological conditions within cells were not considered in these studies. Perturbation of kinase activity in living cells through drug treatment (16, 17) or knocking down/out a specific kinase (18, 19) allows us to monitor consequent changes in the phosphorylation level *in vivo*. However, these approaches would also indirectly affect downstream kinases, which would interfere with the identification of direct substrates (20). To overcome these issues, we and others have employed *in vitro* substrate or sequence motif information, in addition to conducting kinase-perturbed phosphoproteomics using living cells. These approaches have identified endogenous substrate candidates of protein kinase A (PKA), spleen tyrosine kinase (Syk) and Abelson tyrosine kinase (ABL) (21–23).

Although the combined use of *in vitro* substrate information and kinase-perturbed phosphoproteomic profiling has identified putative substrates in some cases, this strategy cannot distinguish substrates of downstream kinases that phosphorylate motifs similar to the target kinase. Therefore, we reasoned that an additional layer of information on kinase-protein interactions including transient interactors should allow more confident identification of endogenous substrates. Indeed, computational analyses have demonstrated that using protein-protein interaction-derived networks and kinase-specific phosphorylation sequence motifs improves the prediction of the substrate specificity (24–26). However, those studies relied on public protein interaction databases in which protein interactions were measured using different cell types in different biological contexts. Furthermore, protein interactions were determined through classical approaches such as affinity-purification mass spectrometry (AP-MS) and yeast two-hybrid studies, which generally cannot capture transient interactors. To determine the endogenous substrates of a target kinase, interactome analysis, which captures not only stable interactions but also transient interactions, should be performed in parallel with kinase-perturbed phosphoproteomic profiling using the same sample.

Recently, proximity labeling approaches, such as proximity-dependent biotin identification (BioID) (27) and an engineered ascorbate peroxidase (APEX) (28), have been developed to capture transient interactions, including kinase-substrate interactions (29–31). BioID is based on biotinylation of proteins proximal (∼10 nm) to a mutant biotin ligase (BirA*)-fused protein of interest; the biotinylated proteins are then captured and identified by means of streptavidin pulldown and LC/MS/MS. Thus, BioID is suitable for interactome analysis to globally capture transient interactions.

In this study, we combined BioID-based interactome analysis with kinase-perturbed phosphoproteomic profiling and substrate motif analysis to establish a general framework for the systematic analysis of endogenous kinase-substrate relationships.

## Experimental procedures

### Cell culture

HEK293T cells were provided by RIKEN BRC through the National BioResource Project of the MEXT/AMED, Japan. HEK293T cells and HeLa cells (HSRRB, Osaka, Japan) were cultured in DMEM (Fujifilm Wako, Osaka, Japan) containing 10% fetal bovine serum (Thermo Fisher Scientific, Waltham, MA) and 100 µg/mL penicillin/streptomycin (Fujifilm Wako).

### Cloning of BirA*-kinase expression vectors

pDEST-pcDNA5-BirA*-FLAG N-term was a gift from Dr. Anne-Claude Gingras (Lunenfeld-Tanenbaum Research Institute at Mount Sinai Hospital, Toronto, Canada). The entry clones pENTR221 CK2A1 and pENTR221 PRKACA were purchased from DNAFORM and RIKEN BRC through the National BioResource Project of the MEXT/AMED, Japan, respectively. CK2A1 or PRKACA coding sequences were cloned into the destination vector pDEST-pcDNA5-BirA*-FLAG N-term with LR clonase 2 (Thermo Fisher Scientific) using the Gateway system. All construct sequences were confirmed by Sanger sequencing (Genewiz, South Plainfield, NJ)

### BioID

For kinase interactome experiments, HEK293T cells in a 10 cm dish were transfected with 40 µL 1.0 mg/mL polyethylenimine (Polysciences, Warrington, PA) and 15 µg plasmid and incubated for 24 h. As the negative control, HEK293T cells in a 10 cm dish were transfected with 40 µL 1.0 mg/mL polyethylenimine for 24 h. All experiments were performed in triplicate. The cells were then incubated for 24 h in culture medium containing 50 µM biotin (Fujifilm Wako), washed, and harvested with ice-cold PBS.

### Drug treatment

HEK293T cells were treated with DMSO, 10 µM CX-4945 (ApexBio, Houston, TX) or 50 µM forskolin (Fujifilm Wako) for 1 hr. Biological triplicates were performed.

### Sample preparation for biotinylated protein identification

Cells were washed and harvested with ice-cold PBS. The proteins were extracted into RIPA buffer (50 mM Tris-HCl (pH 7.2), 150 mM, NaCl, 1% NP-40, 1 mM EDTA, 1 mM EGTA, 0.1% SDS, protease inhibitors cocktail (Sigma-Aldrich, St. Louis, MO), and 1% sodium deoxycholate), and rotated with 300 µg of streptavidin magnetic beads (Thermo Fisher Scientific) for 3 hr at 4 °C. After incubation, the beads were washed with RIPA buffer three times and 50 mM ammonium bicarbonate buffer 3 times, then suspended in 200 µL of 50 mM ammonium bicarbonate buffer. The captured proteins were reduced with 10 mM DTT for 30 min, alkylated with 50 mM iodoacetamide for 30 min in the dark, and digested with Lys-C (w/w 1:100) for 3 h, followed by trypsin digestion (w/w 1:100) overnight at 37 °C, on the beads. The peptides were desalted using StageTip (32) with SDB-XC Empore disk membranes (GL Sciences, Tokyo, Japan) and suspended in the loading buffer (0.5% TFA and 4% acetonitrile (ACN) for subsequent LC/MS/MS analyses.

### Sample preparation for phosphoproteome analysis

Cells were washed and harvested with ice-cold PBS. The proteins were extracted with phase transfer surfactant (33) using a lysis buffer (12 mM sodium deoxycholate (Fujifilm Wako), 12 mM sodium N-lauroylsarcosinate (Fujifilm Wako), 100 mM Tris-HCl (pH 9.0), containing protein phosphatase inhibitor cocktail 1 and 2 (Sigma-Aldrich) and protease inhibitors (Sigma-Aldrich). Protein amount was determined with a BCA protein assay kit and the proteins were reduced with 10 mM DTT for 30 min, and then alkylated with 50 mM iodoacetamide for 30 min in the dark. After reduction and alkylation, proteins were digested with Lys-C (w/w 1:100) for 3 h, followed by trypsin digestion (w/w 1:100) overnight at 37 °C. Then, the peptides were desalted using SDB-XC StageTip.

Phosphopeptides were enriched from 100 µg of tryptic peptides by means of TiO2-based hydroxy acid modified metal oxide chromatography (HAMMOC) (34) and eluted with 0.5% piperidine. Phosphopeptides were labeled with Tandem Mass Tag (TMT) (Thermo Fisher Scientific), desalted using SDB-XC StageTips, fractionated at basic pH (32) and suspended in the loading buffer (0.5% TFA and 4% ACN) for subsequent LC/MS/MS analyses.

### *In vitro* kinase assay using cell extracts

Cells were washed and harvested with ice-cold PBS. Proteins were extracted from HeLa cells with phase transfer surfactant, and the buffer was replaced with 40 mM Tris-HCl (pH7.5) by ultrafiltration using an Amicon Ultra 10K at 14,000 g and 4 °C. Protein amount was confirmed with a BCA protein assay kit and the solution was divided into aliquots containing 100 µg. As described above, proteins were dephosphorylated with TSAP, reacted with recombinant kinase and digested with Lys-C (w/w 1:100) and trypsin (w/w 1:100), using the reported methods (14). Then, phosphopeptides were enriched from the tryptic peptides with HAMMOC, desalted using SDB-XC StageTips and suspended in the loading buffer (0.5% TFA and 4% ACN) for subsequent LC/MS/MS analyses.

### *In vitro* kinase assay using synthetic peptides

A mixture of synthetic peptides (SynPeptide, Shanghai, China) (10 pmol each) was reacted with 0.5 µg of each recombinant kinase (Carna Biosciences, Kobe, Japan) in 100 µL of kinase reaction buffer (40 mM Tris-HCl pH 7.5, 20 mM MgCl2, 1 mM ATP) at 37°C for 3 h. The reaction was quenched by adding 10 µL 10% TFA. Then, the peptides were desalted using SDB-XC StageTip and suspended in the loading buffer (0.5% TFA and 4% ACN) for subsequent LC/MS/MS analyses.

### NanoLC-MS/MS analyses

NanoLC/MS/MS analyses were performed on an Orbitrap Fusion Lumos (Thermo Fisher Scientific), or a Q Exactive (Thermo Fisher Scientific) only for synthetic peptide samples, connected to an Ultimate 3000 pump (Thermo Fisher Scientific) and an HTC-PAL autosampler (CTC Analytics, Zwingen, Switzerland). Peptides were separated on a self-pulled needle column (150 mm length x 100 µm ID, 6 µm opening) packed with Reprosil-C18 AQ 3 µm reversed-phase material (Dr. Maisch, Ammerbuch, Germany). The flow rate was set to 500 nL/min. The mobile phase consisted of (A) 0.5% acetic acid and (B) 0.5% acetic acid in 80% acetonitrile. Three-step linear gradients of 5-10% B in 5 min, 10-40% B in 60 min (for short gradient) or 100 min (for long gradient), and 40-100% B in 5 min were employed.

For BioID samples and phosphopeptides obtained by *in vitro* kinase reaction, the MS scan range was *m/z* 300-1500. MS scans were performed by the Orbitrap with *r* =120,000 and subsequent MS/MS scans were performed by the Orbitrap with *r* = 1,5000. Auto gain control for MS was set to 4.00 × 10^5^ and that for MS/MS was set to 5.00 × 10^4^. The HCD was set to 30.

For TMT-labeled samples, synchronous precursor selection-MS3 (SPS-MS3) (35) was performed. The MS scan range was *m/z* 375-1500. MS scans were performed by the Orbitrap with *r* =120,000, MS/MS scans were performed by the Ion Trap in Turbo mode and MS3 scans were performed by the Orbitrap with *r* = 1,5000. Auto gain control for MS was set to 4.00 × 10^5^, that for MS/MS was set to 1.00 × 10^4^ and that for MS3 was set to 5.00 × 10^4^. The CID was set to 35.

For synthetic peptide samples, the MS scan range was *m/z* 300-1500. MS scans were performed by the Orbitrap with *r* =70,000 and subsequent MS/MS scans were performed by the Orbitrap with *r* = 17,500. Auto gain control for MS was set to 3.00 × 10^6^ and that for MS/MS was set to 1.00 × 10^5^. The HCD was set to 27.

### Database searching

For BioID experiments, the raw MS data files were analyzed by MaxQuant v1.6.2.3 (36). Peptides and proteins were identified by means of automated database searching using Andromeda against the human SwissProt Database (version 2017-04, 20,199 protein entries) with a precursor mass tolerance of 20 ppm for first search and 4.5 ppm for main search and a fragment ion mass tolerance of 20 ppm. Cysteine carbamidomethylation was set as a fixed modification. Methionine oxidation and acetyl on protein N-term were set as variable modifications. The search results were filtered with FDR < 1% at the peptide spectrum match (PSM) and protein levels.

For kinase perturbation experiments, the raw MS data files were analyzed by ProteoWizard (37), ProteomeDiscoverer v1.3.0.339 and MaxQuant v1.6.2.10 to create peak lists based on the recorded fragmentation spectra. Peptides and proteins were identified by means of automated database searching using Mascot v2.4 (Matrix Science, London) against the human SwissProt database (version 2017-04, 20,199 protein entries) with a precursor mass tolerance of 5 ppm and a fragment ion mass tolerance of 20 ppm. Cysteine carbamidomethylation and TMT tags on lysine and peptide N-termini were set as fixed modifications. Methionine oxidation and phosphorylation on serine, threonine and tyrosine were set as variable modifications. The search results were filtered with FDR < 1% at the peptide level. As described previously (21), we also used the additional criteria that at least three successive y- or b-ions with a further two or more y-, b- and/or precursor-origin neutral loss ions were observed, based on the error-tolerant peptide sequence tag concept (38). Phosphosite localization was confirmed with a site-determining ion combination method (39). This method is based on the presence of site-determining y- or b-ions in the peak lists of the fragment ions, which unambiguously identify the phosphosites. Only singly phosphorylated peptides with confident phosphosites were accepted for the following analysis. Site-confirmed phosphopeptides were remapped to protein sequences for protein annotation. Peptides assigned to multiple proteins in the SwissProt database were rejected.

For *in vitro* kinase reaction using HeLa cells, the raw MS data files were analyzed by ProteoWizard to create peak lists based on the recorded fragmentation spectra. Peptides and proteins were identified by means of automated database searching using Mascot v2.4 (Matrix Science, London) against the human SwissProt database (version 2017-04, 20,199 protein entries) with a precursor mass tolerance of 5 ppm and a fragment ion mass tolerance of 20 ppm. Cysteine carbamidomethylation was set as a fixed modification. Methionine oxidation and phosphorylation on serine, threonine and tyrosine was set as variable modifications. The search results were filtered with FDR < 1% at the peptide level. We also used the additional criteria for the identification and site-confirmation of phosphopeptides same as kinase-perturbed experiments.

### Data analysis

For BioID samples, the peak area of each peptide in MS1 was quantified using MaxQuant. Missing values were imputed with values representing a normal distribution around the detection limit of the mass spectrometer. A two-tailed Welch’s t-test was performed comparing the BirA*-kinase group to the control group. For the following analysis, out of Majority protein IDs, the lead protein was used.

For TMT samples, the peak intensities of reporter ions were obtained from MS3 scan. Missing values were imputed with values representing a normal distribution around the detection limit of the mass spectrometer. The ratio of drug-treated to control was logged (base 2) for each phosphopeptide, and averaged to each phosphosite.

For synthetic peptides, phosphorylation ratio was calculated as follows:

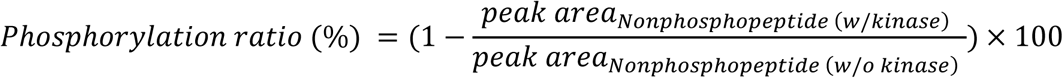

The protein-protein interaction (PPI) network analysis was performed with STRING version 11.0 (40) and the medium-confidence PPI network (score = 0.4) based on the experimental evidences was used. GO analysis was performed with the Database for Annotation, Visualization, and Integrated Discovery (DAVID) version 6.8 (41, 42). The background was set as *Homo sapiens*.

### Motif score

The probability of observing residue *x* in position *i* is computed as follows:

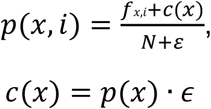

where ***f***_*x*,i_ is the frequency of observing residue ***x*** at position ***i*** and N is the total number of sequences. ***c(x*)** is a pseudo count function which is computed as the probability of observing residue b in the proteome, *p*(*x*), multiplied by ɛ, defined as the square root of the total number of sequences used to train the PWM. This avoids infinite values when computing logarithms. Probabilities are then converted to weights as follows:

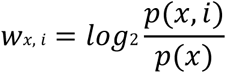

where *p*(*x*)*=* background probability of amino acid ***x; p(x, i***) = corrected probability of amino acid ***x*** in position ***i***; **W*x*, i** *=* PWM value of amino acid ***x*** in position ***i***. Given a sequence *q* of length *l*, a score *λ* is *then* computed by summing log_2_ weights:

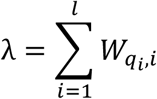

where *qi* si the *ith* residue of *q*. In this study, the score was computed using the flanking ±7 residues surrounding each phosphosite.

## Results

### Defining kinase-interacting proteins using BioID

To identify kinase-interacting proteins, we first performed BioID experiments (Fig. 1A). We selected CK2 and PKA as target kinases, because their functions, localizations and putative substrates have been relatively well investigated (8). BirA*-fused kinases (BirAk) were individually transfected into HEK293T cells, and non-transfected cells were used as a negative control. We observed a broad spectrum of biotinylated proteins by streptavidin blotting for the BirAk-expressing cells, while no signals were detectable for the control cells (Fig. S1A), confirming the validity of our BioID experiments. After the transfection and biotinylation, biotinylated proteins were enriched with streptavidin beads and digested to obtain tryptic peptides, which were analyzed by nanoLC/MS/MS and quantified using label-free quantification (43). As a result, 2,218 proteins and 1,795 proteins were quantified in at least two of the three replicates in the CK2- and PKA-expressing cells, respectively. Among them, 574 and 518 proteins were considered as interacting proteins of CK2 and PKA, respectively, after applying the cut-off values (Fig. 1B, Table S1). Importantly, CSNK2B and PRKAR1A, which form heteromeric complexes with the corresponding kinases, were significantly enriched as kinase interactors (Fig. 1B), suggesting that the tagged kinases form the heteromeric complexes *in vivo* and are likely to function in the same manner as the endogenous kinases.

**Figure 1.**
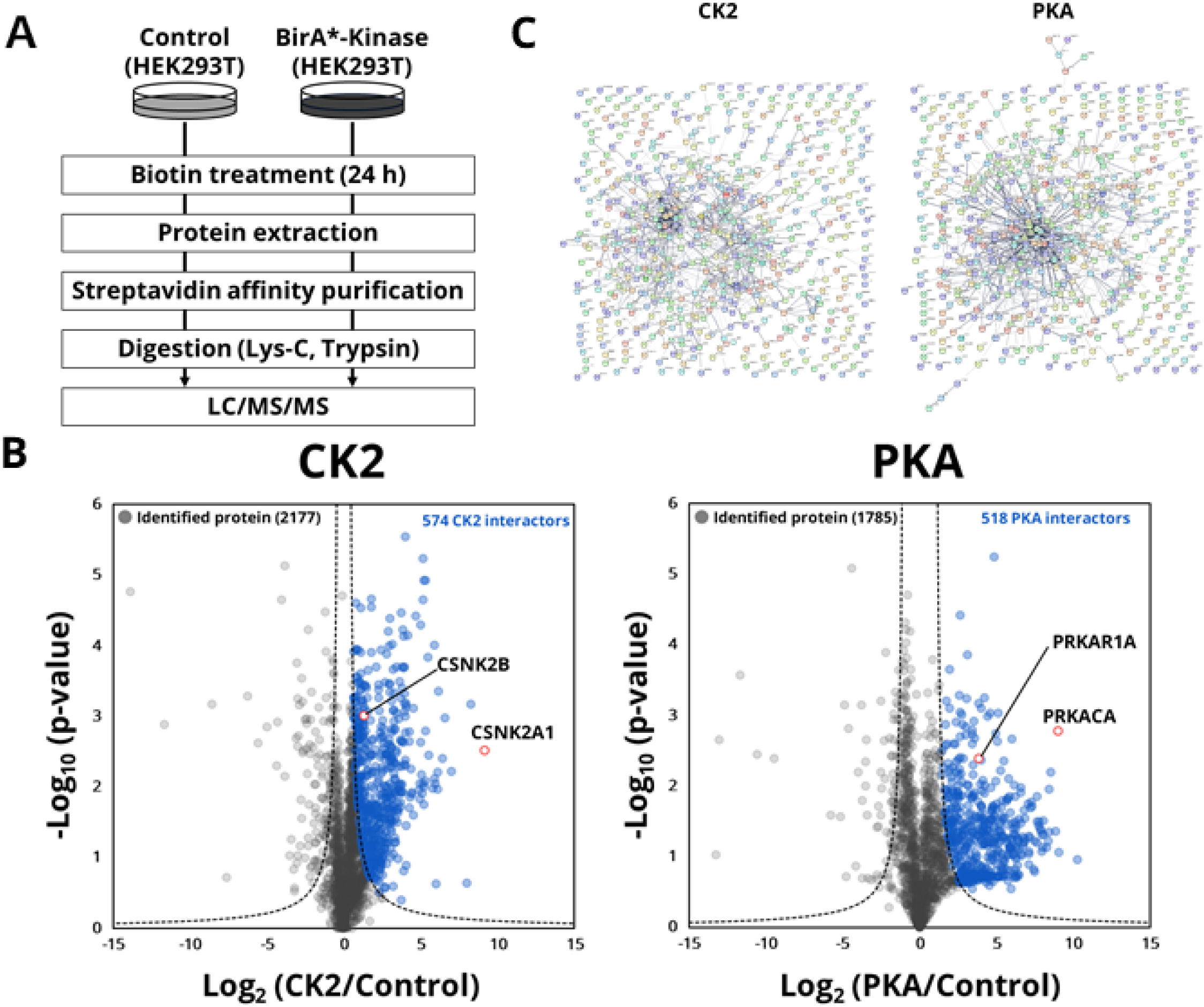
Identification of kinase-interacting proteins with BioID. (A) Workflow to identify kinase-interacting proteins with BioID. (B) The ratio of BirAk to control (log_2_ (BirAk/control)) and the negative value of log_10_ *p*-values (Welch’s t-test) are plotted for each protein. Proteins were filtered by cutoff lines (*y= c/(x – x*_*0*_ *=* curvature, *x*_*0*_ *=* minimum fold change) as previously described (44). Cutoff parameters for CK2 were *c =* 1 and *x*_*0*_ *=* 0.3, and those for PKA were *c =* 1 and *x*_*0*_ *=* 1. (C) PPI network of interacting proteins.

To benchmark our method, we evaluated the biotinylation of previously known interactors for CK2 and PKA (40). We found that the known interactors exhibited significantly higher ratios of BirAk/control compared to all proteins quantified in this study (Fig. S1B). Then, we mapped CK2- or PKA-interacting proteins (n = 574 and n = 518, respectively) on the STRING (40)-based protein interaction networks. We found that 395 proteins (corresponding to 69%, p-value < 10^−16^.) and 371 proteins (72%, p-value < 10^−16^) composed single large networks (that is, any given pair of proteins is connected directly and/or indirectly) (Fig. 1C). This indicates that our BioID experiments capture biologically meaningful and proximal proteins to the target kinases; the remaining approximately 30% of the interactors are likely to be transient interactors that would be difficult to detect by other methods. Finally, we performed Gene Ontology (GO) enrichment analyses for the sets of kinase-interacting proteins (Fig. S1C). We found that GO terms related to “cellular proliferation” and “apoptosis” were enriched in the CK2-interacting proteins, consistent with the known functionality and biology of the CK2 family (45). For the PKA-interacting proteins, we observed the term “cell-cell adhesion”, in line with PKA’s known function (46).

Altogether, the results suggest that our data have a high level of confidence, and represent a rich source of CK2 and PKA interactomes including both stable and transient interactors.

### Phosphoproteomic profiling of responses to kinase perturbations

Having established the CK2 and PKA interactomes, we next sought to perform kinase-perturbed phosphoproteome profiling to identify phosphosites regulated by the target kinases (Fig. 2A). To this end, HEK293T cells were treated with either an ATP-competitive CK2 inhibitor, CX-4945, also known as silmitasertib (47) or a PKA activator, forskolin (48). Following protein extraction and tryptic digestion, phosphopeptides were enriched by means of titania chromatography (34), labeled with TMT reagents and analyzed by nanoLC/MS/MS. To simplify the interpretation of kinase-phosphosite relationships, we only considered mono-phosphopeptides in subsequent analyses; as a result, 9,313 and 9,140 mono-phosphosites were quantified in triplicate from CX-4945- and forskolin-treated cells, respectively (Table S2). Down-regulation of known CK2 substrates such as TOP2A (S1377), LIG1 (S66) and MRE11 (S649) and up-regulation of known PKA substrates such as CAD (S1406), NDE1 (S306) and FLNA (S2152) were confirmed, supporting the validity of our results (Fig. S2A). Furthermore, expected sequence motifs for acidophilic CK2 and basophilic PKA were clearly enriched in perturbed phosphosites (2-fold change by CK2 inhibitor or PKA activator with p-value < 0.05) (Fig. S2B), demonstrating that the data obtained in this experiment are meaningful and of high quality.

**Figure 2.**
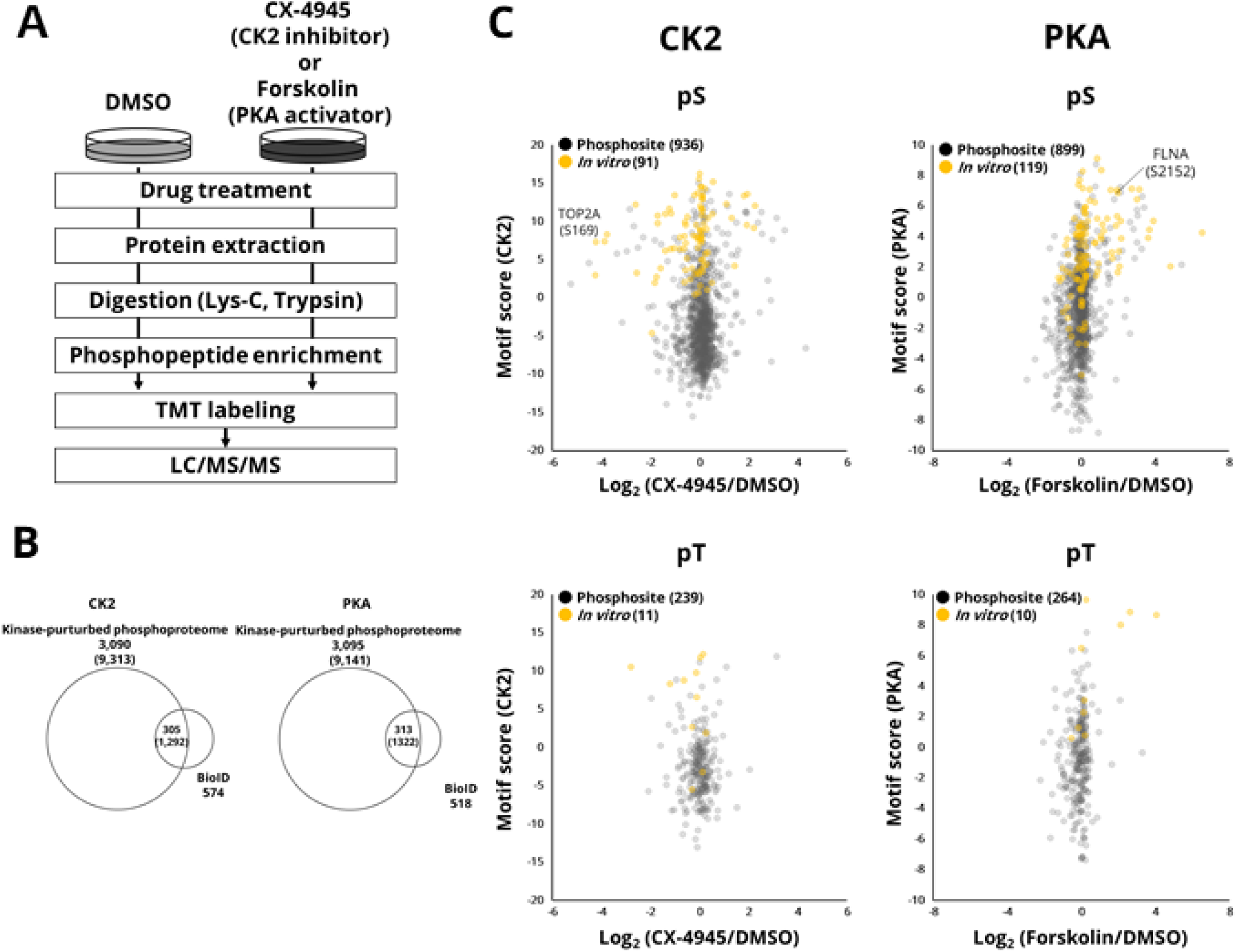
Identification of phosphosites regulated by the target kinase. (A) Quantitative phosphoproteomics to profile phosphorylation changes in the presence of target kinase inhibitor/activator. (B) Overlap of identified proteins between phosphoproteomics and the target kinase interactors. Numbers in parentheses are the numbers of phosphosites. (C) The distribution of phosphorylation ratios and motif scores for phosphosites identified in cells treated with inhibitor or activator. Yellow color indicates sites of *in vitro* substrates.

To combine the results obtained from the phosphoproteomic and BioID experiments, we mapped the identified phosphorylation sites onto the kinase-interacting proteins; 1,292 and 1,322 phosphosites were attributed to the 305 and 313 proteins interacting with CK2 and PKA, respectively (Fig. 2B). We then set the cut-off, 2-fold change on average (n=3) with p-value < 0.05, to further select the regulated phosphosites, resulting in the identification of 67 and 74 phosphosites for CK2 and PKA substrates, respectively.

### Sequence motif analysis

Our kinase-perturbed phosphoproteomic analysis identified all regulated sites, which could include sites indirectly regulated due to effects such as perturbation of downstream kinases of the target kinases or off-target effects of the drugs used for perturbation. Thus, the next step is to select phosphosites that match the consensus phosphorylation motif of the kinase, which has been shown to be a useful predictor in kinase identification (21).

Position weight matrices (PWMs) (49) represent the signatures of the sequences flanking the phosphosites targeted by a given kinase (Fig. S3). Here columns in the matrix represent relative positions from phosphosites, while rows represent residues. The values in the matrix are log2-transformed probabilities of a residue’s occurrence at a position. These probabilities are used to predict putative substrates (see methods). This approach has been implemented in several kinase-substrate prediction methods and applied to *in vivo* kinase–substrate discovery. We built the PWM based on thousands of substrates identified in our previous study (14) and this study (Fig. 2C), and calculated the motif scores for the substrate candidates. Finally, 33 CK2 substrates and 52 PKA substrates met the criteria (motif score > 2, Table S3). These substrates include well-known substrates such as TOP2A (S1377) for CK2 and FLNA (S2152) for PKA. Note that only two proteins (NAP1L1, TOP2A) are annotated as CK2 interactors in STRING, while none of them are described as PKA interactors in STRING, meaning that transient interactors captured by BioID might have met the above criteria. The GO enrichment analysis revealed that RNA splicing-related proteins were enriched as CK2 substrates, which is consistent with CK2 being an RNA splicing-related kinase (50), while cell adhesion-related molecules were identified as PKA substrates, in accordance with PKA’s involvement in the cell-cell adhesion pathway (46).

### Missense mutations near phosphosites of CK2 and PKA substrates

Having discovered putative direct substrates of CK2 and PKA, we next sought to assess the relationship between the kinase and substrates in the context of cancer biology (51, 52). The effect of amino acid substitutions in substrates on the kinase-substrate relationship has been analyzed using kinase substrate sequence specificity (53, 54). Information on amino acid substitutions that occur within seven residues around the phosphosites in Table S3 was extracted from the cancer genomics database cBioPortal (55, 56). As a result, 121 missense mutations from 33 CK2 substrates and 189 missense mutations from 52 PKA substrates were found. For substrate sequences containing these mutations, we used PWM scores to predict phosphorylation preference by kinases and compared them to wild-type sequences (Fig. S4). Then, wild-type/mutant substrate pairs (five substrate pairs for CK2 and one pair for PKA) that showed potential loss of phosphorylation by mutation were selected for *in vitro* kinase assays using synthetic peptides (Fig. 3). As a result, mutation-induced reduction in phosphorylation stoichiometry was observed for 5 out of the 6 pairs. These results indicate that missense mutations near phosphosites of substrates negatively affect phosphorylation by CK2 or PKA, and thus would be expected to affect downstream signaling networks in cancer cells bearing these mutations.

**Figure 3.**
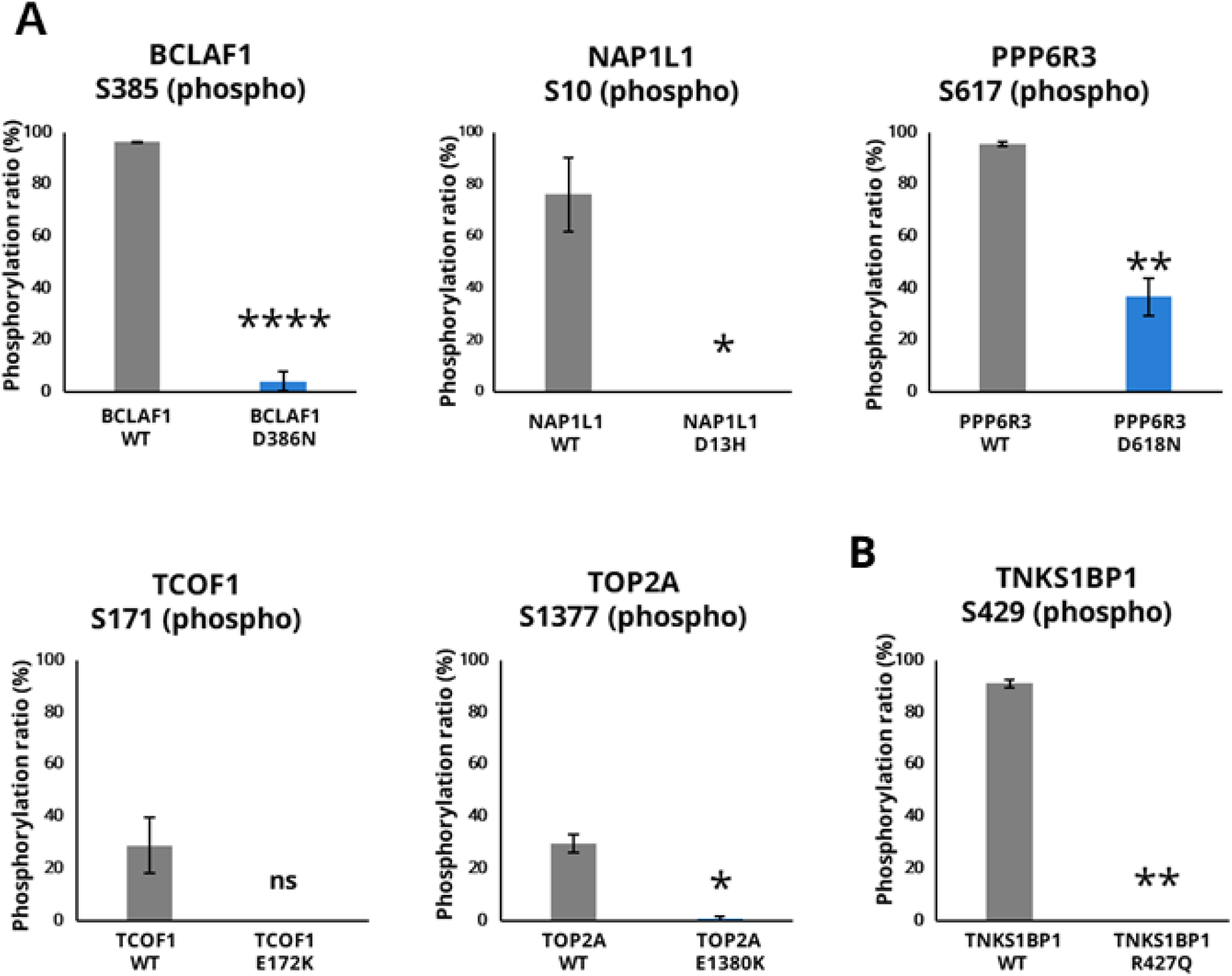
*In vitro* kinase assay of putative kinase substrates using synthetic peptides. (A) CK2 substrates (B) PKA substrate. Significance levels are indicated as follows,****; *p*-value < 0.0001, **; *p*-value < 0.01, *; *p*-value < 0.05 and ns; *p*-value > 0.05.

## Discussion

Knowledge of kinase-substrate relationships is critical to understand intracellular signaling networks. We believe the present study is the first to identify endogenous kinase substrates by combining three approaches: proximity labeling, kinase-perturbed phosphoproteomics, and motif preference analysis. Using this strategy, we were able to identify novel endogenous substrates as well as known endogenous substrates. This combined approach offers many advantages. First, BioID is able to identify transient interactors that could not be found by conventional methods. Second, motif preference analysis can be applied to any sequence, as it scores matches against substrate motifs rather than against substrates obtained in *in vitro* kinase assay. Finally, this combined approach (BioID, kinase-perturbed phosphoproteomics, and motif analysis) should be applicable to any kinase as a universal method.

One of the applications of this approach is the identification of endogenous kinase substrate mutations associated with cancer. For example, our present analysis demonstrates the potential impact of known oncogenic mutations on phosphorylation-related signaling pathways, and illustrates the usefulness of motif analysis for screening. We believe the methodology described here will be useful for large-scale discovery of endogenous kinase substrates.

## Data Availability

The mass spectrometry proteomics data have been deposited at the ProteomeXchange Consortium (http://proteomecentral.proteomexchange.org) via the jPOST partner repository (http://jpostdb.org) (57) with the data set identifier PXD019664.

## Acknowledgments

We would like to thank Dr. Anne-Claude Gingras for providing BirA* constructs and members of the Department of Molecular & Cellular BioAnalysis for fruitful discussions. This work was supported by JST Strategic Basic Research Program CREST (No. 18070870), AMED Advanced Research and Development Programs for Medical Innovation CREST (18068699) and JSPS Grants-in-Aid for Scientific Research No. 17H03605 to Y.I., No. 20H03241 to K.I. and 20H04845 to N.S.

## Author contributions

T.N, K.I., N.S. and Y.I. designed the research. T.N. performed the research and analyzed data. T.N, K.I. and Y.I. wrote the paper.

## Abbreviations

ACN: Acetonitrile
AP-MS: Affinity-purification mass spectrometry
BioID: Proximity-dependent biotin identification
BirA: Biotin ligase
CK2: Casein kinase 2
GO: Gene Ontology
HAMMOC: Hydroxy acid modified metal oxide chromatography
LC/MS/MS: Liquid chromatography/tandem mass spectrometry
PKA: Protein kinase A
PPI: Protein-protein interaction
PSM: Peptide spectrum match
PWMs: Position weight matrices
TMT: Tandem Mass Tag

